# Mechanistic Elucidation of the Antitumor Properties of a Novel Death Receptor 5 Activator

**DOI:** 10.1101/700906

**Authors:** Mengxiong Wang, Mary E. Law, Bradley J. Davis, Elham Yaaghubi, Amanda F. Ghilardi, Renan B. Ferreira, Chi-Wu Chiang, Olga A. Guryanova, Daniel Kopinke, Coy D. Heldermon, Ronald K. Castellano, Brian K. Law

## Abstract

Disulfide bond Disrupting Agents (DDAs) are a new chemical class of agents recently shown to have activity against breast tumors in animal models. However, it is unknown how DDAs trigger cancer cell death without affecting nontransformed cells. As demonstrated here, DDAs are the first compounds identified that upregulate the TRAIL receptor DR5 through both transcriptional and posttranscriptional mechanisms. At the protein level, DDAs alter DR5 disulfide bonding to increase steady-state DR5 levels and oligomerization, leading to downstream Caspase 8 and 3 activation. DDAs and TRAIL synergize to kill cancer cells and are cytotoxic to HER2+ cancer cells with acquired resistance to the EGFR/HER2 tyrosine kinase inhibitor. Investigation of the mechanisms responsible for DDA selectivity for cancer cells reveals that DDA-induced upregulation of DR5 is enhanced in the context of EGFR overexpression, and DDA-induced cytotoxicity is strongly amplified by MYC overexpression. Together, the results demonstrate selective DDA lethality against oncogene-transformed cells, DDA-mediated DR5 upregulation and protein stabilization, and DDAs against drug-resistant and metastatic cancer cells. DDAs thus represent a new therapeutic approach to cancer therapy.

## Introduction

Breast cancer remains a major cause of cancer mortality despite improvements in the survival of patients resulting from earlier detection and an increasing number of molecularly targeted therapeutic agents. The primary events associated with breast cancer deaths are tumor acquisition of resistance to currently available drugs and the development of metastatic disease. Endocrine therapies are a mainstay for Estrogen Receptor and Progesterone Receptor positive tumors, and include Estrogen Receptor antagonists and modulators, as well as Aromatase inhibitors that block estrogen synthesis. Cdk4/6 and mTORC1 inhibitors are FDA approved for hormone receptor-positive tumors that have acquired resistance to endocrine therapies. Monoclonal antibodies and tyrosine kinase inhibitors are available for tumors overexpressing Human Epidermal growth factor Receptor 2 (HER2) or Epidermal Growth Factor Receptor (EGFR) tyrosine kinases. HER2+ cancers, however, frequently acquire resistance to targeted agents, and HER2-specific drugs have limited efficacy unless combined with cytotoxic chemotherapy agents.

Tumors that lack expression of Estrogen Receptor, Progesterone Receptor, and HER2 are termed “Triple-Negative” Breast Cancers (TNBCs). In general, TNBCs lack sensitivity to targeted therapies, with the exception of Poly-ADP Ribose Polymerase (PARP) inhibitors that have efficacy against BRCA1/2-mutant cancers. EGFR is overexpressed in a substantial fraction of TNBCs and has been proposed as a therapeutic target (Costa, Shah et al., 2017), but so far, EGFR-targeted therapies have not exhibited sufficient activity against EGFR+ TNBCs to merit FDA approval. EGFR and HER2 specific drugs are designed to act through the mechanism of oncogene addiction. A weakness of this approach is that cancers can easily acquire redundant (horizontal) or downstream (vertical) alterations that permit escape from addiction to the founder oncogene. In contrast, synthetic lethality, as exemplified by the use of PARP inhibitors against BRCA1/2-mutant cancers, relies on drug actions that are manifested selectively in the context of tumor-associated genetic alterations. Given the current limitations of EGFR and HER2 targeted drugs in treating breast cancer, new agents that act through a synthetic lethal mechanism rather than by oncogene addiction could benefit patients with EGFR+ TNBCs and drug resistant HER2+ tumors.

Previous reports (Ferreira, Law et al., 2015, Ferreira, Wang et al., 2017, Law, Ferreira et al., 2016) describe a novel chemical class of anticancer agents termed Disulfide bond Disrupting Agents (DDAs). DDAs selectively kill cancer cells that overexpress EGFR or HER2 (Ferreira et al., 2015), and DDA treatment is associated with decreased expression of EGFR, HER2, and HER3 and reduced phosphorylation of the pro-survival serine/threonine kinase Akt. DDA-mediated cancer cell death is also associated with activation of the Unfolded Protein Response (UPR) (Ferreira et al., 2017). The Endoplasmic Reticulum (ER) stress that triggers the UPR can activate multiple cell death pathways (reviewed in (Wang, Law et al., 2018)). However, the specific cell death mechanisms engaged by the DDAs remain unexplored.

Irremediable ER stress can drive apoptosis by upregulating the Death Receptor 5 (DR5) protein through transcriptional mechanisms (Abdelrahim, Newman et al., 2006, Tiwary, Yu et al., 2010, Xu, Su et al., 2012, Yamaguchi & Wang, 2004). Activation of DR5 by its ligand, TNF-Related Apoptosis-Inducing Ligand (TRAIL), is well known for selectively killing cancer cells without effects on nontransformed cells, and for manageable side effects in clinical trials (Ashkenazi & Dixit, 1999, Griffith, Wiley et al., 1999, Rieger, Naumann et al., 1998). However, TRAIL and other DR5 agonists have not met expectations in clinical trials, in part because cancer cells can easily become TRAIL-resistant by downregulating DR5 (Cheong, Lee et al., 2011, James, Seibel et al., 2015, Jazirehi, Kurdistani et al., 2014). A therapeutic strategy for increasing DR5 levels and activating downstream DR5 apoptotic signaling could bypass the resistance to TRAIL and DR5 agonist antibodies often observed in the clinic.

Herein we describe novel mechanisms by which DDAs drive cancer cell death by upregulating the TRAIL receptor DR5. DDAs increase DR5 expression through transcriptional mechanisms and post-transcriptional means that involve increased DR5 steady-state protein levels and disulfide-bond mediated DR5 oligomerization. Additionally, oncogene-dependent mechanisms responsible for the selective killing of cancer cells by DDAs without toxicity to normal tissues are demonstrated.

## Results

### DDAs activate extrinsic apoptosis pathway to kill EGFR+ and HER2+ cancers

Previous studies revealed that breast cancer cells that overexpress EGFR or HER2 are particularly sensitive to DDAs (Ferreira et al., 2015, Ferreira et al., 2017, Wang, Ferreira et al., 2019). This work employed MDA-MB-468 TNBC cells and BT474 luminal B cells as models of EGFR and HER2 overexpressing breast cancer, respectively. Consistent with these previous studies, DDAs show extensive anticancer effects *in vivo*. Mice bearing orthotopic xenograft BT474 tumors were treated with Vehicle (DMSO) or different doses of our most highly potent DDA tcyDTDO (Wang et al., 2019) for 5 days. Representative hematoxylin & eosin (H&E) histochemical staining of tumor tissue revealed that the tumors were substantially necrotic after tcyDTDO treatment (Fig. 1A, left panels). The corresponding Cleaved Caspase 3 staining confirmed that the tumor cell death was due to caspase-dependent apoptosis (Fig. 1A, right panels). To investigate which pathway is important for modulating DDA-induced tumor cell death, we screened multiple cell death axes. Immunoblot analyses showed that the DDA tcyDTDO (Fig. 1B) significantly increased the expression level of DR5 (Fig. 1C, left panel), suggesting that DDAs activate the extrinsic apoptotic pathway in cancer cells. T47D cells were used as a model for low EGFR/HER2 luminal A cancers as employed previously (Ferreira et al., 2015, Ferreira et al., 2017, Wang et al., 2019). These studies showed that T47D cells engineered to overexpress EGFR or HER2 are highly sensitive to DDAs. Therefore, we examined whether DR5 expression increases in EGFR overexpressing T47D cells treated with DDAs. TcyDTDO only activated ER stress and upregulated DR5 in T47D/EGFR cancer cells, but not T47D/vector cells (Fig. 1C, right panel). The combination of tcyDTDO and Cycloheximide (CHX), a protein synthesis inhibitor, reversed DDA induced upregulation of ER stress and DR5, indicating that DDA-triggered ER stress is responsible for increasing DR5 levels. To further validate that DR5 is critical for regulating the sensitivity of cancer cells to DDAs, we employed shRNA to generate DR5 knock down MDA-MB-468 lines (Fig. 1D). Knockdown of DR5 decreased PARP cleavage and partially reduced sensitivity of cancer cells to tcyDTDO treatments as measured by MTT assays (Fig. 1D).

**Figure 1.**
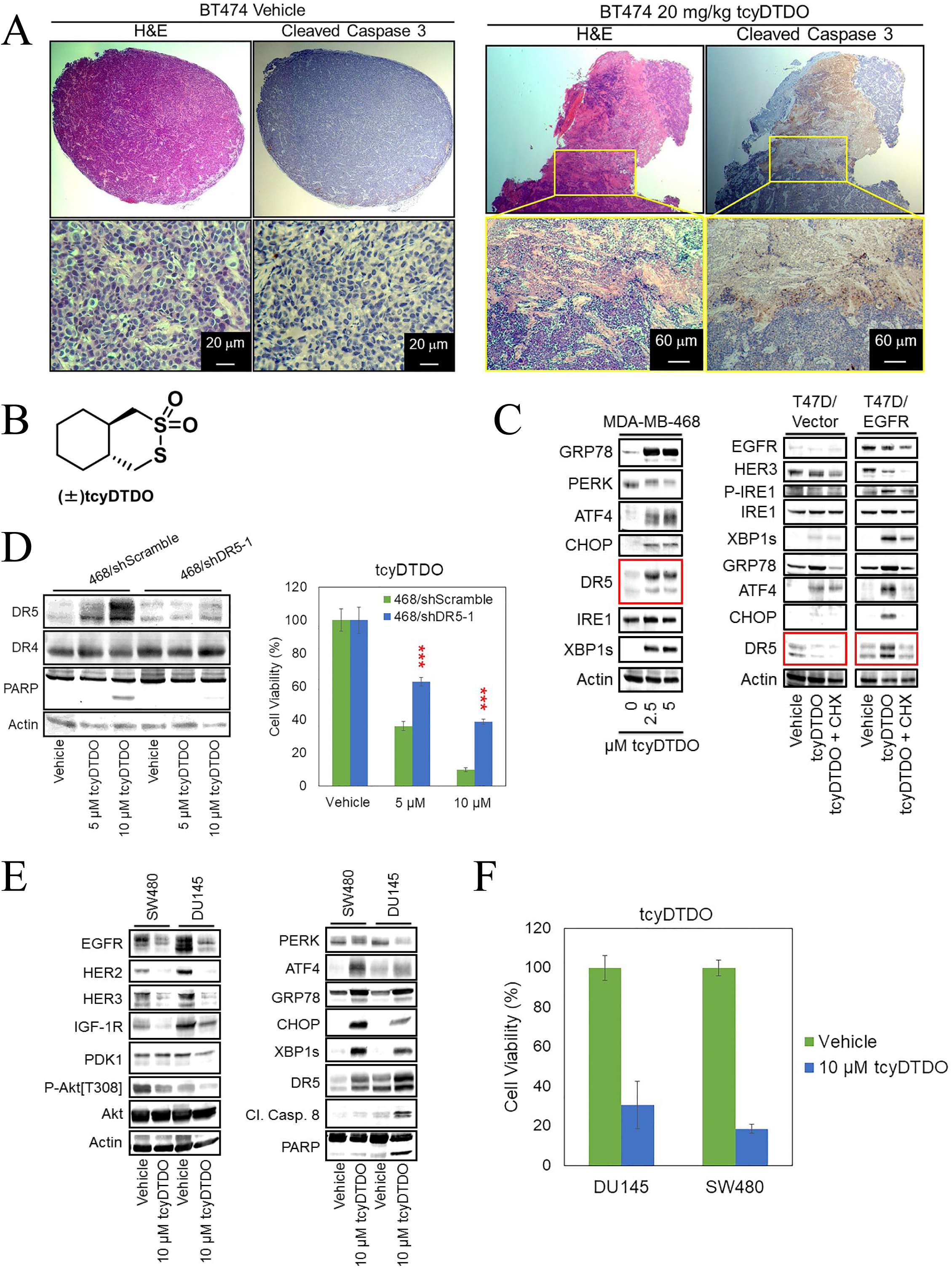
DDAs activate extrinsic apoptosis pathway to kill EGFR+ and HER2+ cancers. A. Tumor bearing mice were separated into two groups of three mice each and treated once daily for five days with either vehicle (DMSO) or 20 mg/kg tcyDTDO. Mice were sacrificed on day five (3 h after treatments) and tumor samples were collected for Hematoxylin and Eosin (H&E) staining and Cleaved Caspase 3 staining. B. Structure of DDA tcyDTDO. C. MDA-MB-468 and the indicated T47D stable cell lines were treated for 24 h has indicated and cell extracts were analyzed by immunoblot. D. Left panel. Immunoblot analyses were performed on the indicated MDA-MB-468 lentivirally transduced shRNA knock-down cell lines with 5 or 10 µM tcyDTDO treatments. Right panel. MTT assays were performed on the same cell lines treated with the indicated concentrations of tcyDTDO for 72 h. E. SW480 and DU145 cell lines were treated as indicated for 24 h and analyzed by immunoblot. F. MTT assays were performed on SW480 and DU145 cells with indicated treatments for 72 h. Data information: Results are presented as the mean ± standard deviation of five replicate determinations. Significance was determined using Student’s t-test with P ≤ 0.001 (***).

In previous studies, we demonstrated that DDA treatment kills various types of cancer cells that overexpress EGFR and/or HER2 (Wang et al., 2019). The human tumor lines DU145 (prostate) and SW480 (colon) are two such representatives of DDA-sensitive non-breast cancer cell lines. It is shown here that 10 µM tcyDTDO downregulated the expression levels of EGFR/HER2/HER3, induced Akt dephosphorylation, and induced ER stress in these cells (Fig. 1E). Importantly, tcyDTDO also triggered the extrinsic apoptosis pathway as measured by the upregulation of DR5, CC8 and PARP cleavage, and significantly reduced cell viability (Fig. 1F). These data suggest that DDAs activate the extrinsic apoptosis pathway to kill EGFR+ and HER2+ cancers.

### DDAs upregulate DR5 through both transcriptional and posttranscriptional mechanisms, by altering DR5 disulfide bonding pattern and promoting its oligomerization

Studies have demonstrated that ER stress increases the transcription of the gene encoding DR5 through activation of transcription factors ATF4 and CHOP (Chen, Hu et al., 2016, Kim, Hong et al., 2013, Liu, Shi et al., 2016). Therefore, we examined whether tcyDTDO activates a previously described(Cheong et al., 2011) DR5 transcriptional reporter construct. TcyDTDO (5 µM) stimulated a several-fold increase in the activity of the DR5 reporter (Fig. S1A), suggesting a transcriptional mechanism. To validate this finding in cancer cells, time-course treatments (2-16 h) of MDA-MB-468 cells with 5 µM tcyDTDO were performed, and the mRNA levels of DR5 were measured by RT PCR and quantitative PCR. As shown in Fig. 2A and S1B, the increase of DR5 mRNA level was only detected at 2 h after tcyDTDO treatment, which only induced a 1.5 fold change. Since CHOP is important in upregulating the transcription of DR5, we constructed stable MDA-MB-468 and SKBR3 cell lines encoding either tetracycline-inducible control vector (468/tet-Puro) or CHOP (468/tet-CHOP), and stably knocked down CHOP in BT474 cells to study the effects of CHOP on DR5 levels in response to DDAs. Results indicate that CHOP overexpression failed to increase DR5 levels with or without DDA treatment (Fig. S1C, D), and CHOP knockdown only partly blocked DDA-mediated DR5 upregulation (Fig. S1E). These data suggest that tcyDTDO only slightly increased the transcription of DR5. ER stress also reduces protein synthesis through PERK-dependent phosphorylation of eIF2α so we examined the effect of tcyDTDO on protein synthesis. TcyDTDO suppressed protein synthesis in a concentration-dependent manner similarly to the protein synthesis inhibitor Cycloheximide (CHX) (Fig. 2B). Therefore, we hypothesized that DDAs may regulate DR5 levels through posttranscriptional and posttranslational mechanisms as well as through transcriptional means. We constructed stable MDA-MB-468 cell lines to inducibly express the long alternative splice variant of DR5 (468/tet-DR5), or Firefly Luciferase (468/tet-fLuc) as a control, to study the effects of DDAs on DR5 steady state protein levels. In the 468/tet-DR5 cells, doxycycline + tcyDTDO robustly upregulated DR5 and this was associated with strong induction of Caspase 8 and 3 cleavage (Fig. 2C). In contrast, in the 468/tet-fLuc cells, doxycycline + tcyDTDO induced a weaker upregulation of DR5 and Caspase 3 and 8 cleavage. Examining the effects of all possible combinations of doxycycline, tcyDTDO, and TRAIL on 468/tet-DR5 cells demonstrated that the highest levels of the long form of DR5 were achieved by combining doxycycline and tcyDTDO (Fig. 2D), suggesting that tcyDTDO may stabilize DR5 at the protein level. These treatments did not alter the levels of the closely related DR5 homolog DR4 (Fig. 2E). Comparison of tcyDTDO with other activators of the UPR demonstrated that even at similar levels of ER stress induction, tcyDTDO produced the highest level of steady-state DR5 (Fig. 2E). However, it is interesting to note that levels of DR4 were increased by ER stressors that disrupt Ca^2+^ homeostasis (Thapsigargin), reduce disulfide bonds (dithiothreitol (DTT)), and inhibit proline *cis/trans*-isomerization (Cyclosporine A (CsA)), while DR4 levels were not significantly altered by tcyDTDO. The DR5 extracellular domain contains seven disulfide bonds(Cha, Sung et al., 2000, Mongkolsapaya, Grimes et al., 1999) and DR5_Long_ contains an additional unpaired Cys, Cys 209, that is not present in DR5_Short_ (Fig. 2F, upper panel). Given our previous proposal that DDA actions are mediated through effects on disulfide bond formation(Ferreira et al., 2015), we considered that tcyDTDO might alter the disulfide-bonding pattern of DR5. Immunoblot analysis of 468/tet-DR5 samples, which were prepared under reducing or non-reducing conditions in the presence of 100 mM *N*-ethylmaleimide (NEM) to prevent disulfide exchange after lysis, was used to compare effects of multiple ER stress inducers on DR5 (Fig. 2F, lower panel). Analysis of the samples under reducing conditions revealed that Thapsigargin, Tunicamycin, and tcyDTDO, induced comparable ER stress as measured by GRP78, but tcyDTDO had the strongest effect in upregulating DR5. Analysis of the samples under non-reducing conditions indicated that only tcyDTDO treatment induced migration of DR5_Long_ monomer and increased accumulation of the disulfide-bonded multimeric form of DR5 at the top of the gel. Moreover, studies have shown that MG132 and b-AP15 induce DR5 accumulation by inhibiting the proteasome and proteasome-associated deubiquitinases, respectively (Cheong et al., 2011, Chiu, Lin et al., 2008, Hetschko, Voss et al., 2008, Oh, Deng et al., 2017). Multiple replicate experiments demonstrated that 5 µM tcyDTDO, but not Thapsigargin, Tunicamycin, b-AP15, MG132, or TRAIL, reduced the electrophoretic ability of both DR5_Long_ and DR5_Short_ under non-reducing conditions in various cell lines (Fig. S2A-C). Together, these results suggest that the DDA tcyDTDO altered the disulfide bond pattern of DR5_long_, and increased the formation of DR5 multimers, by promoting the formation of inter-disulfide bonds.

**Figure 2.**
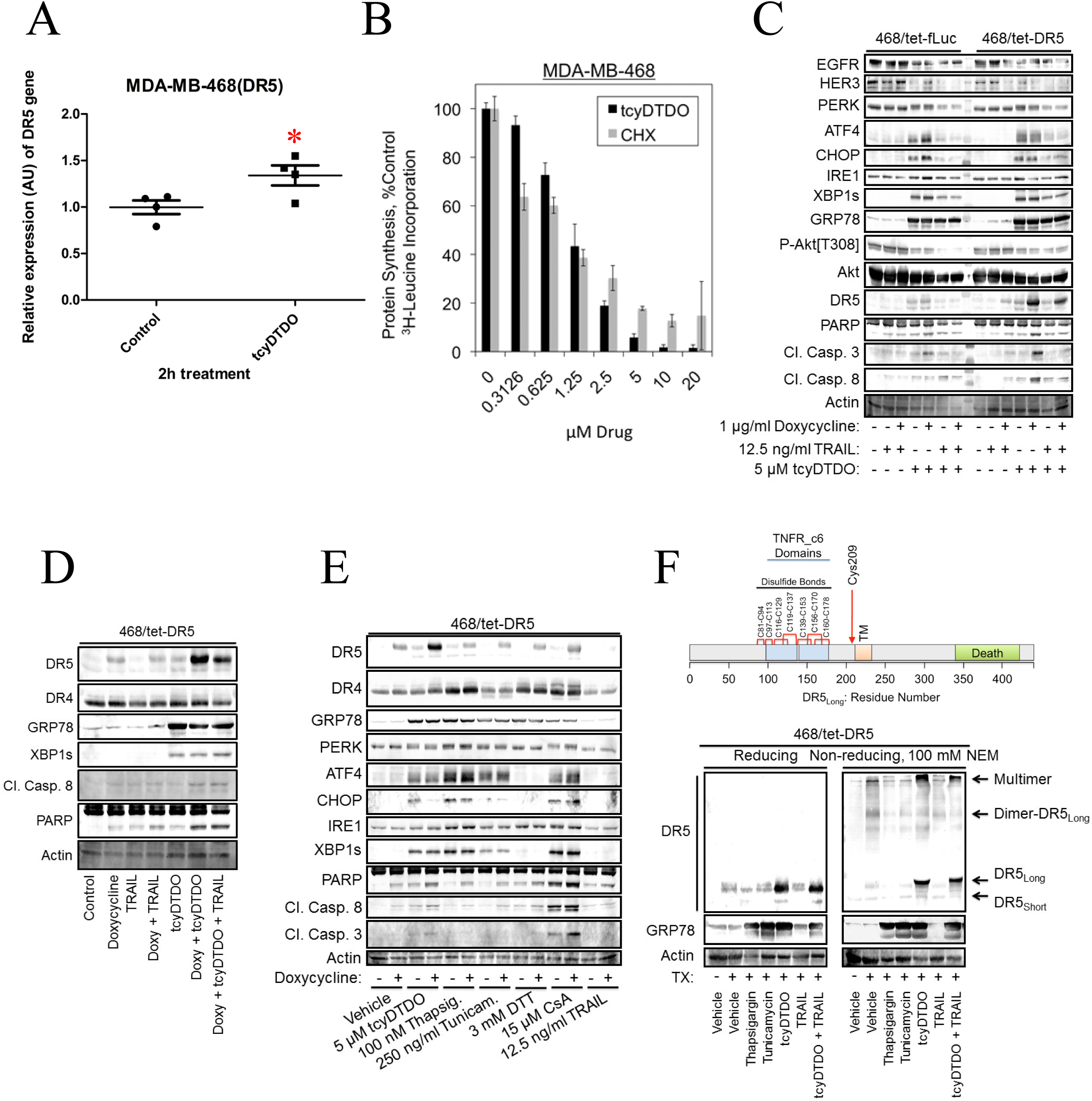
DDAs upregulate DR5 through transcriptional and post-transcriptional mechanisms. A. MDA-MB-468 cells were treated with DMSO or 5 µM tcyDTDO for 2 h. Total RNA was isolated from each sample and converted to cDNA using reverse transcriptase. The mRNA levels of DR5 were measured using real-time qPCR, and relative mRNA expression was calculated. Fold change was calculated by normalizing all values to the untreated group. T-test showed p= 0.0395. B. Protein synthesis was assessed by ^3^H-leucine incorporation in MDA-MB-468 cells treated with the indicated concentrations of tcyDTDO or cycloheximide (CHX). C. Immunoblot analysis of MDA-MB-468 cells engineered to express DR5 in a tetracycline-inducible manner (468/tet-DR5) and the corresponding control cell line (468/tet-fLuc) after the indicated 24 h treatments. D. Immunoblot analysis of 468/tet-DR5 cells treated separately or with combinations of 1 µg/ml Doxycycline, 12.5 ng/ml TRAIL, and 5 µM tcyDTDO for 24 h. E. Immunoblot analysis of 468/tet-DR5 cells treated with the indicated agents for 24 h. F. Upper panel. Diagram based on DR5 crystal structures showing the presence of seven intramolecular disulfide bonds and an unpaired Cysteine residue in the extracellular domain of DR5. Lower panel. Immunoblot analysis of DR5 from 468/tet-DR5 cells treated for 24 h as indicated under non-reducing and reducing conditions. Cell extraction in the presence of N-ethylmaleimide NEM (100 mM) was used to limit thiol-disulfide exchange under non-reducing conditions.

### DDA upregulation of DR5 correlates with sensitization to TRAIL

As we demonstrated that DDAs are able to stabilize DR5 protein level through a post-transcriptional mechanism, we wanted to investigate whether DDA can sensitize cancer cells to the DR5 ligand TRAIL. Cell viability studies were performed in MDA-MB-468 (Fig. 3A) and BT474 (Fig. 3B) cells and the results were analyzed using the Chou-Talalay method (Chou & Talalay, 1981, Chou & Talalay, 1984) to determine whether tcyDTDO and TRAIL synergize to induce cell death. Combination Indices (CIs) less than one indicate synergy. The results showed that the combination of tcyDTDO and TRAIL induced strong synergy in killing both cell lines across different concentrations. CRISPR/Cas9-mediated genome editing technology was applied to generate DR5 knockout BT474 clones that were compared with a polyclonal population of empty vector control cells (Fig. 3C, left panel). The cell viability assay confirmed that ablation of DR5 expression significantly reduced sensitivity of cells to tcyDTDO treatment, or combined tcyDTDO/TRAIL treatment (Fig. 3C, right panel). It is noted that the complete deletion of DR5 expression did not fully rescue cell death from tcyDTDO + TRAIL treatments, so we hypothesized that DR4 might be activated by tcyDTDO after DR5 was knocked out. Vector control cells and DR5 knockout clones from different cell lines were treated in the presence or absence of tcyDTDO and cell lysates were prepared in reducing or non-reducing conditions in the presence of 100 mM NEM (Fig. 3D). The results indicated that tcyDTDO did not upregulate the total protein levels of DR4, but strongly induced DR4 oligomerization in DR5 knock out cells (Fig. 3D), which supports this hypothesis. Moreover, tcyDTDO more strongly stimulated DR5 oligomerization in T47D/EGFR cells than in T47D/Vector, suggesting that EGFR overexpression potentiates the effects of tcyDTDO on DR5 (Fig. 3E). Together, the results in Fig. 3 demonstrate that the DDA tcyDTDO sensitizes cancer cells to TRAIL treatment by inducing DR5 upregulation and oligomerization.

**Figure 3.**
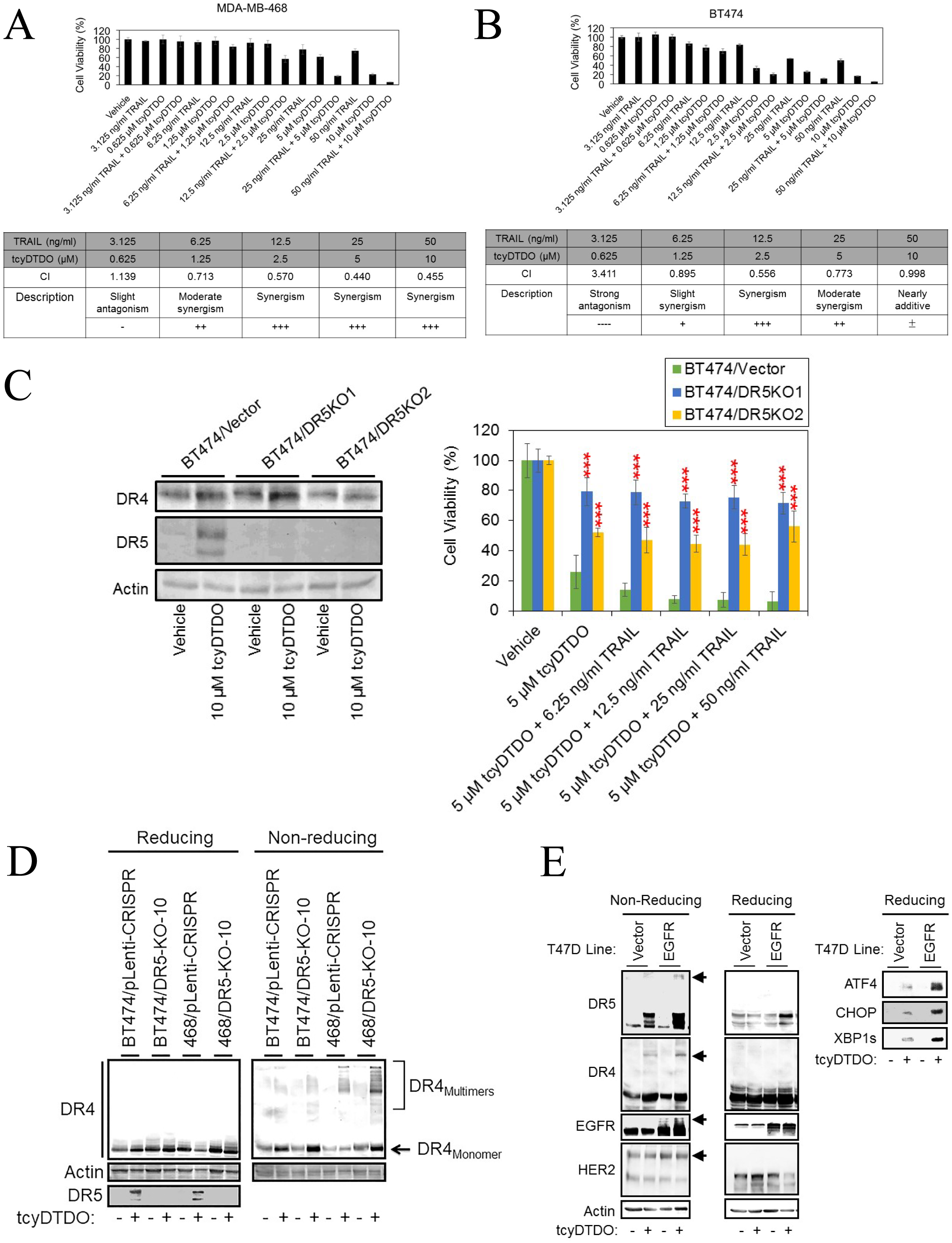
DDA upregulation of DR5 correlates with sensitization to TRAIL. A. B. MTT cell viability assays performed on MDA-MB-468 and BT474 cells, respectively, after 72 h of the indicated treatments (upper panel) and analyzed using the Chou-Talalay method to calculate combination indices (CIs) (lower panel). [CIs: = 1, > 1, and < 1 represent additivity, antagonism, and synergy, respectively.] Graphs represent the average of quadruplicate determinations ± standard deviation. C. Left panel. Immunoblot analysis of BT474 control and DR5 knockout (DR5-KO) clones after a 24 h treatment with vehicle or 10 µM tcyDTDO. Right panel. MTT cell survival assays performed on BT474 control and DR5 knockout clones after 72 h of the indicated treatments. D. Immunoblot analysis showing comparison of DR4 oligomerization in MDA-MB-468 and BT474 vector control or DR5 knockout clones. E. The indicated T47D cell lines were treated in the presence or absence of 5 µM tcyDTDO for 24 h. Samples were collected under reducing and non-reducing conditions for immunoblot analyses.

### DDAs kill Lapatinib-resistant breast cancer cells

It is widely accepted that most cancer deaths are due to tumor resistance to available drugs, and to metastasis. We previously isolated a cell line termed HCI-012 (Wang et al., 2019) isolated from a Patient-Derived Xenograft (PDX) model of the same name. The HCI-012 xenograft originated from a patient with HER2+ breast cancer that metastasized and resisted treatment with cytotoxic and targeted (Lapatinib/Trastuzumab) therapy (DeRose, Wang et al., 2011). We selected sublines of the HCI-012 cells to model drug-resistant and metastatic breast cancers in two steps. We first isolated HCI-012 sublines capable of colonizing mouse lungs and liver after injection into either the tail vein or mammary gland (Fig. 4A). Second, continuous growth of the HCI-012 cells from liver metastases (012/LVM) in the presence of either 5 µM or 10 µM Lapatinib produced the Lapatinib resistant sublines 012/LVM/LR5 and 012/LVM/LR10, respectively. Immunoblot analysis of these lines showed that the Lapatinib resistant lines expressed higher levels of EGFR, HER2, and DR5 (Fig. 4B). As expected, Lapatinib cytotoxicity was reduced in the Lapatinib-resistant lines (Fig. 4C), but the sensitivity of all three cell lines to tcyDTDO was similar, demonstrating the lack of cross-resistance between Lapatinib and tcyDTDO. Treatment of the control cell lines with Lapatinib increased expression of EGFR and HER2, and this effect was more dramatic in the Lapatinib resistant lines (Fig. 4D). These observations suggest that the Lapatinib resistance observed may be due to Lapatinib upregulating EGFR and HER2 in addition to higher basal EGFR and HER2 levels in these cells. TcyDTDO partially decreased EGFR expression and Akt phosphorylation in control and Lapatinib-resistant cells. To study anticancer activities of tcyDTDO against Lapatinib resistant tumors *in vivo*, eight mice were injected with HCI012/LVM/LR10 cells in their mammary fat pads. After tumor volumes of the primary tumors reached over 100 mm^3^, mice were treated once daily with either Vehicle or 20 mg/kg of tcyDTDO for five consecutive days by intraperitoneal injection. Primary and metastatic tumor samples were collected for H&E staining and Cleaved Caspase 3 staining. Representative H&E staining of tumor tissue showed that both the primary and metastatic tumors from tcyDTDO treated mice were extensively necrotic, and large numbers of dying cells were observed surrounding the necrotic areas (Fig. 4E). Cleaved Caspase 3 staining of serial sections confirmed that the apoptotic pathway was activated, especially in and surrounding the tumor necrotic area (Fig. 4E). Importantly, tcyDTDO only induced apoptosis of cancer cells, while the normal muscle and stomach tissues remained intact, indicating selectivity of tcyDTDO toward cancer cells (Fig. 4E).

**Figure 4.**
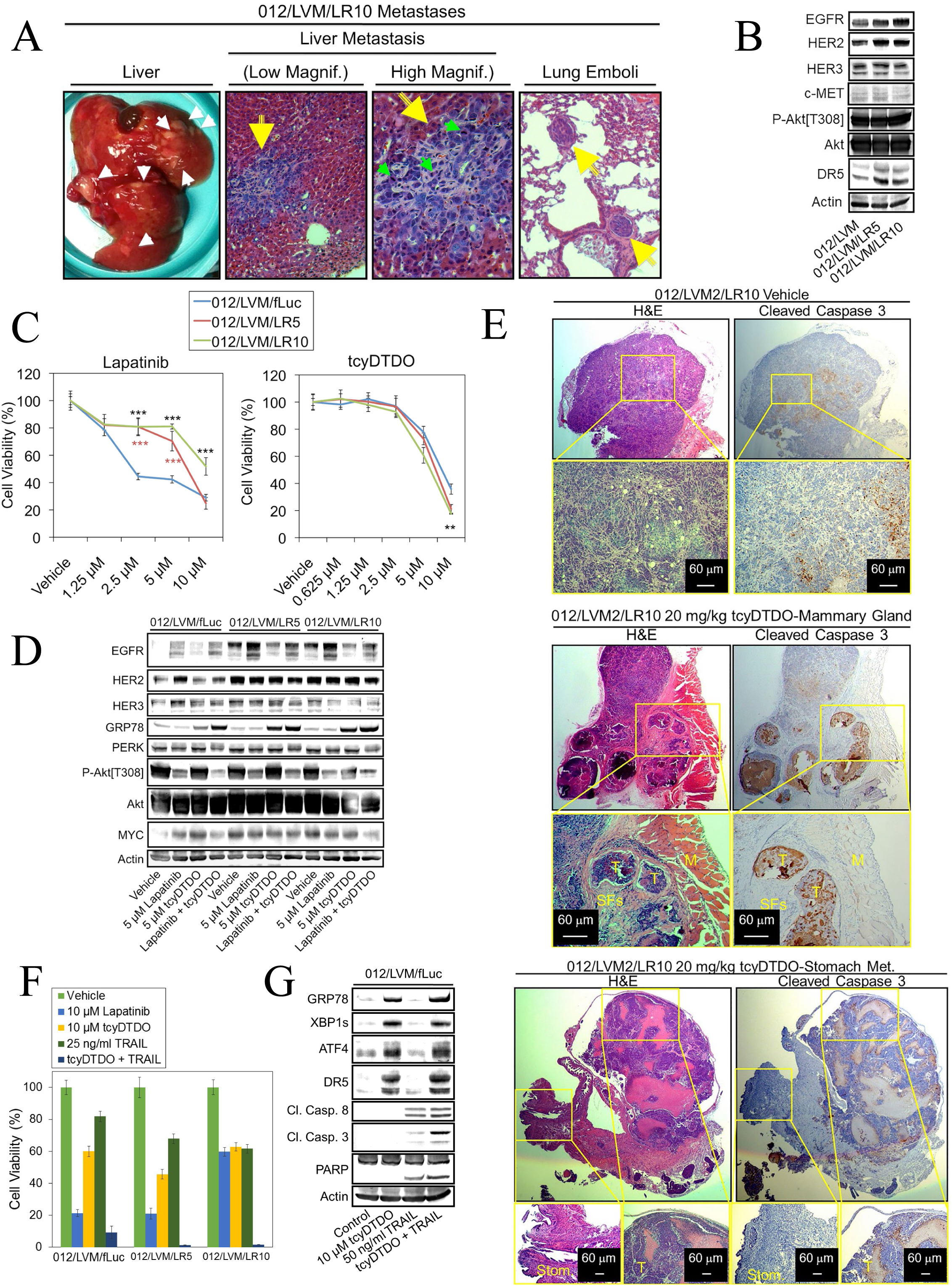
DDAs kill Lapatinib-resistant cancer cells. A. Micrographs of liver metastases and lung tumor emboli generated in mice by an HCI-012 cell line selected for liver metastasis (LVM) and growth in the presence of 10 µM Lapatinib (LR10). B. Immunoblot analysis of parental liver metastatic HCI-012 cells (012/LVM), or sublines selected for growth in 5 (LR5) or 10 µM (LR10) Lapatinib. C. MTT cell viability assays on the indicated cell lines after treatment for 72 h with increasing concentrations of Lapatinib (left panel) or tcyDTDO (right panel). D. Immunoblot analysis of the indicated cell lines treated for 24 h with the specified concentrations of Lapatinib, tcyDTDO, or Lapatinib + tcyDTDO. E. Tumor bearing mice were separated into two groups of four mice each and treated for five days with either vehicle (DMSO) or 20 mg/kg of tcyDTDO. Mice were sacrificed at day five (3 h after treatments) and tumor samples and metastatic lesions were collected for Hematoxylin and Eosin (H&E) staining and Cleaved Caspase 3 staining. Abbreviations used include: M, muscle tissue; T, tumor tissue; SFs, stromal fibroblasts. F. MTT cell viability assays performed after a 72 h treatment with 10 µM Lapatinib, 10 µM tcyDTDO, 25 ng/ml TRAIL, or tcyDTDO + TRAIL. G. Immunoblot analysis of 012/LVM/fLuc cells treated for 24 h as specified. 012/LVM/fLuc cells were labeled using a Lentiviral Firefly Luciferase vector as described in Materials and Methods.

The combination of tcyDTDO and TRAIL is an even better strategy against Lapatinib resistant 012/LVM lines, as the viability of these cells was somewhat decreased by treatment with either tcyDTDO or TRAIL, but tcyDTDO + TRAIL dramatically reduced the viability of all three lines (Fig 4F). TcyDTDO and TRAIL-induced cell death was apparent as demonstrated by reduced cell numbers and apoptotic cell morphology in photomicrographs (Fig. S2D). Consistent with these observations, co-treatment with tcyDTDO and TRAIL resulted in maximal Caspase 8, Caspase 3, and PARP cleavage, indicative of the induction of apoptosis through the extrinsic death pathway (Fig. 4G).

### Caspases mediate DDA/TRAIL-induced downregulation of EGFR and PDK1

As observed with the HCI-012 cell lines, tcyDTDO and TRAIL co-treatment killed MDA-MB-468 and BT474 cells more effectively than either agent alone (Fig. 5A and S2E) did. The pan-Caspase inhibitor Q-VD-OPH partially rescued the cells from tcyDTDO + TRAIL-mediated cell death (Fig. 5A, B). This rescue was associated with reversal of the rounded and apoptotic morphology of cells treated with tcyDTDO to the attached, spread morphology observed with the vehicle treated cells. Immunoblot analysis indicated that tcyDTDO increased expression of DR5, but not DR4, and tcyDTDO + TRAIL strongly decreased EGFR, HER2, HER3, and IGF-1R expression and Akt phosphorylation (Fig. 5C). TcyDTDO + TRAIL appeared to induce the degradation of Akt, PDK1, and PERK based on the loss of the parent bands and formation of more rapidly migrating species. Many of these effects were reversed by Caspase inhibition. EGFR (He, Huang et al., 2006) and Akt (Medina, Afsari et al., 2005) were previously demonstrated to be Caspase substrates. We are not aware of reports showing that PDK1 or PERK are degraded via a Caspase-dependent mechanism. TRAIL + tcyDTDO produced near complete PARP cleavage, which was reversed by Q-VD-OPH. In some systems, ER stress upregulates ATG12 and initiates autophagy (Wang, Kang et al., 2014), or induces necroptosis (Livezey, Huang et al., 2018). In MDA-MB-468 cells, tcyDTDO + TRAIL did not increase expression of the autophagy marker ATG12 or the necroptosis marker phospho-MLKL, suggesting that cell death is predominantly through Caspase-dependent apoptosis (Fig. 5C). Similarly, in 012/LVM/fLuc cells, tcyDTDO + TRAIL strongly decreased EGFR, HER2, HER3, and PDK1 levels and Akt phosphorylation, and increased PARP cleavage (Fig. 5D). These results indicate that DDAs and TRAIL cooperate to downregulate EGFR, HER3, PERK, Akt, and PDK1. Further, this downregulation correlates with the greatest degree of PARP cleavage, and these effects were reversed by Caspase inhibition. In contrast, the ER stress and DR5 upregulation caused by tcyDTDO were not altered by Caspase inhibition. The inability of Q-VD-OPH to overcome the ER stress response and the associated inhibition of protein synthesis and cell proliferation likely explains its inability to restore cell viability to control levels. Together, the results suggest that the model in Fig. 5E is operative in breast cancer cell lines, wherein activation of the TRAIL/DR5 pathway leads to the activation of Caspases that directly execute cell death in part through decreased expression of elements of the HER/PI3K/PDK1/Akt survival pathway.

**Figure 5.**
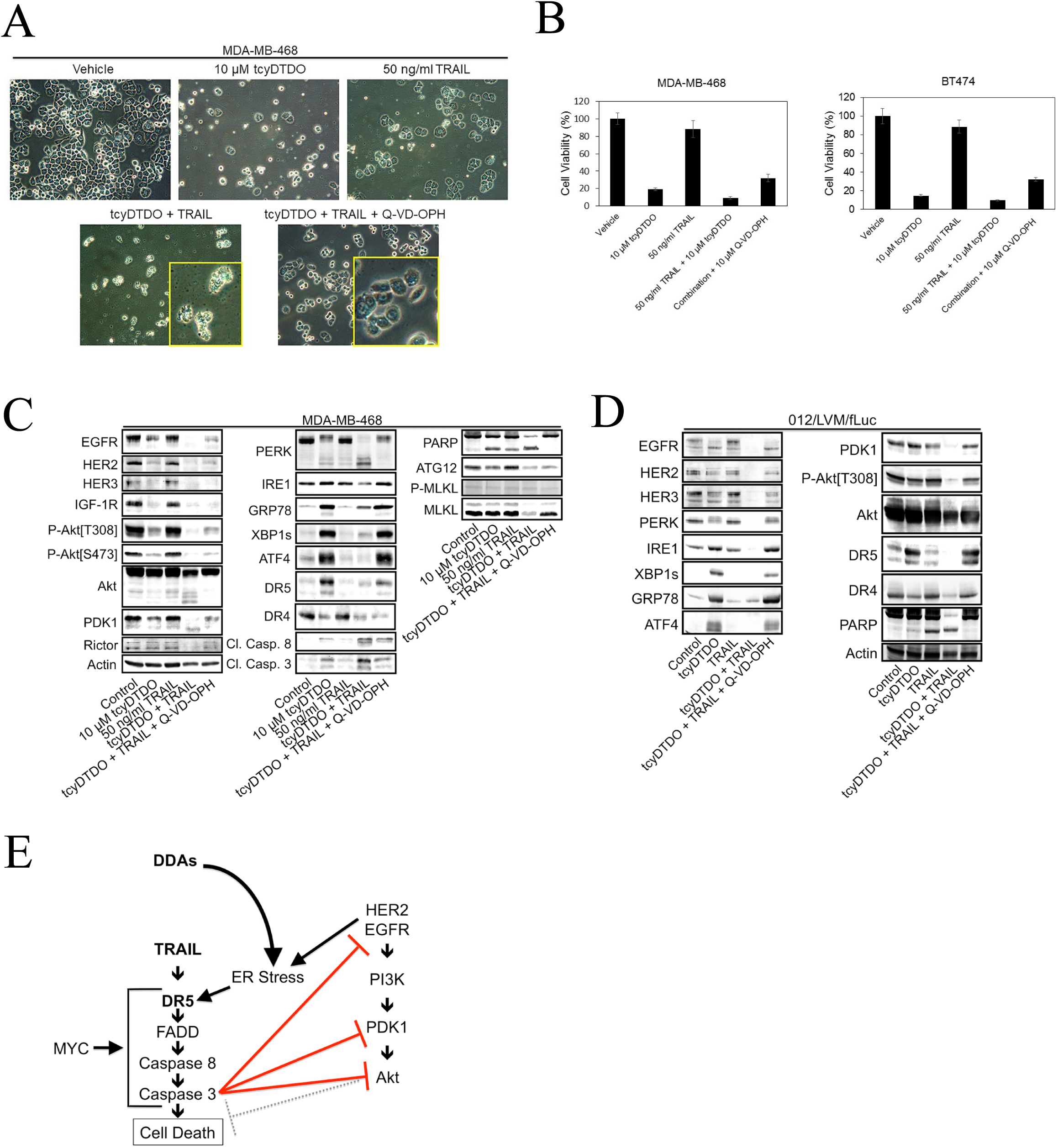
Caspases mediate DDA/TRAIL-induced downregulation of EGFR and PDK1. A. Micrographs of MDA-MB-468 cells treated as indicated for 24 h showing cytotoxicity of tcyDTDO and TRAIL and prevention of cell death by the Caspase inhibitor Q-VD-OPH (10 µM). B. MTT assays were performed on MDA-MB-468 and BT474 cells with indicated treatments for 72 h. C. Immunoblot analysis of MDA-MB-468 cells treated as in A. D. Immunoblot analysis of 012/LVM/fLuc cells treated for 24 h as indicated. E. Model for how DDAs and TRAIL reduce levels of PERK, EGFR, PDK1, and Akt by inducing their caspase-dependent degradation.

### DDAs selectively kill oncogene-transformed cells

Cancer cells that overexpress EGFR or HER2 are particularly sensitive to DDAs (Ashkenazi, Pai et al., 1999, Costa et al., 2017). The cancer lines chosen as representative of EGFR and HER2-overexpression, MDA-MB-468 and BT474, respectively, also show amplification of the locus coding for MYC (Rieger et al., 1998). Breast cancers with MYC and HER2 co-amplification are metastatic and exhibit a tumor initiating cell/cancer stem cell phenotype and poor prognosis (Griffith et al., 1999). Kaplan-Meier analyses demonstrate that breast tumors with high expression of both EGFR and MYC are likewise associated with decreased patient survival (Fig. 6A). To investigate whether overexpression of MYC increases the sensitivity of cancer cells to DDAs, we engineered MCF10A human mammary epithelial cells to ectopically express EGFR, MYC, or both proteins using retroviral vectors. Photomicrographs and immunoblot analysis indicate that neither tcyDTDO nor TRAIL induced changes in cell morphology or PARP cleavage in the vector control cell line (Fig. 6B, C). In contrast, overexpression of EGFR or MYC either separately or together sensitized MCF10A cells to tcyDTDO, TRAIL, or tcyDTDO + TRAIL-induced cell death, but did not sensitize these cells to Lapatinib. The pan-Caspase inhibitor Q-VD-OPH reversed the cell rounding and apoptotic morphology caused by tcyDTDO and tcyDTDO + TRAIL, but did not reverse tcyDTDO-mediated ER stress. Together, the results in Fig.6 indicate that overexpression of either EGFR or MYC is sufficient to sensitize cells to tcyDTDO or TRAIL-mediated cell death. These results are consistent with DDAs and TRAIL selectively killing oncogene-transformed cells through overlapping mechanisms, and suggest that MYC/EGFR may serve as predictive biomarkers of tcyDTDO efficacy.

**Figure 6.**
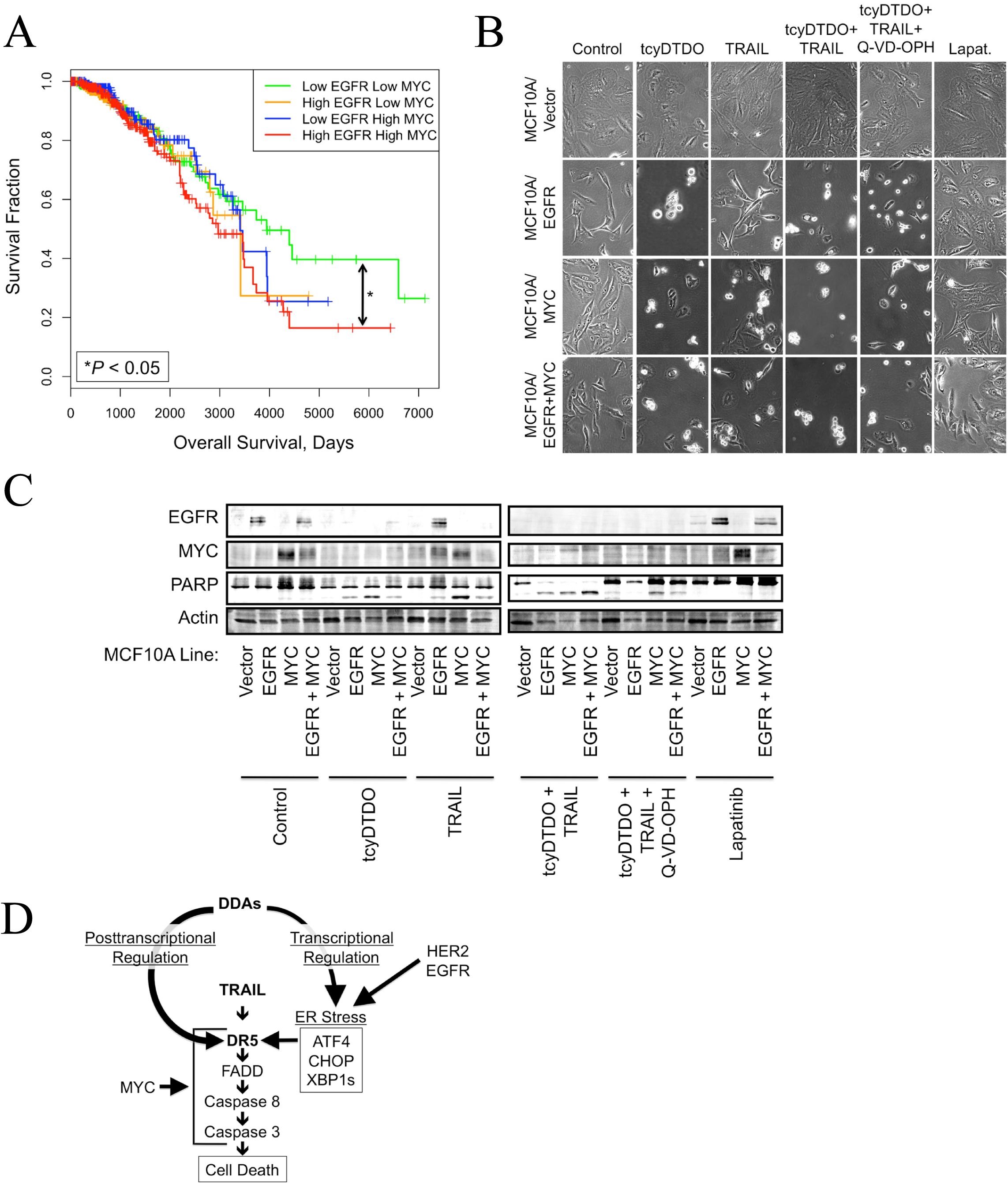
MYC or EGFR overexpression is sufficient to confer DDA cytotoxicity. A. Kaplan-Meier plot showing that patients with tumors overexpressing both EGFR and MYC are associated with a significantly worse survival than are patients with tumors expressing low levels of both EGFR and MYC. Patients ranked based on the expression of EGFR and MYC were classified into four groups, named “low EGFR low MYC (N=359)”, “low EGFR high MYC (N=220)”, “high EGFR low MYC (N=225)”, and “high EGFR high MYC (N=372)”. Overall Survival (OS) was compared among these groups. The F test was used to compare the variance between groups (*P* > 0.05, ns; *P* ≤ 0.05, *; *P* ≤ 0.01, **; *P* ≤ 0.001, ***). B. Micrographs showing the morphology of the indicated MCF10A stable cell lines after 24 h treatment with 5 µM tcyDTDO, 12.5 ng/ml TRAIL, tcyDTDO + TRAIL, tcyDTDO + TRAIL + 10 µM Q-VD-OPH, or 10 µM Lapatinib. C. Immunoblot analysis of stable MCF10A cell lines treated for 24 h as in B. D. Model for how DDAs activate TRAIL/DR5-induced cell death in an oncogene-dependent manner. In the context of EGFR or HER2 overexpression, DDAs elevate ER stress resulting in transcriptional upregulation of DR5. Through a second mechanism, DDAs alter DR5 disulfide bonding to promote DR5 protein stabilization, oligomerization, and activation of pro-apoptotic signaling. Cytotoxicity of DDAs and TRAIL is also potentiated in MYC overexpressing cells.

## Discussion

Great effort has been expended in testing soluble TRAIL and DR5 agonist antibodies as anticancer agents (reviewed in (Kretz, von Karstedt et al., 2018, Yuan, Gajan et al., 2018)). For the most part, these trials have not demonstrated dramatic anticancer activity, which has been attributed to short *in vivo* half-lives and trimer dissociation for soluble TRAIL agonists (Jiang, Kim et al., 2011, Kim, Youn et al., 2011, Seol, Park et al., 2003), and the inability of bivalent agonist antibodies to trigger the same receptor trimerization induced by soluble TRAIL. DR4 and DR5 can be nonfunctional in cancer cells due to decreased expression of the proteins (Song, Szczepanski et al., 2010), constitutive protein internalization (Jin, McDonald et al., 2004, Mert & Sanlioglu, 2017, Zhang, Yoshida et al., 2009), or alterations in posttranslational modifications (Liang, Xu et al., 2018, Micheau, 2018, Yoshida, Shiraishi et al., 2007). Approaches directed toward solving these problems have included the generation of more stable trimeric forms of TRAIL (Han, Moon et al., 2016, Lamanna, Smulski et al., 2013, Liu, Su et al., 2017, Pan, Xie et al., 2013), and the induction of increased DR5 expression through pharmacological agents that induce ER stress to activate *DR5* transcription (Abdelrahim et al., 2006, Tiwary et al., 2010, Xu et al., 2012, Yamaguchi & Wang, 2004, Zhang, Inukai et al., 2011, Zou, Yue et al., 2008). Other strategies have included increasing DR5 half-life by decreasing its proteasomal degradation by inhibiting the proteasome (He, Huang et al., 2004, Hetschko et al., 2008, Hougardy, Reesink-Peters et al., 2008) or proteasome-associated deubiquitinases (DUBs) (Oh et al., 2017). We are not aware of pharmacological approaches that: a) cause DR5 accumulation and oligomerization; and b) stimulate downstream caspase activation and cancer cell death through mechanisms involving altered DR5 disulfide bonding.

Together, our results suggest the model in Fig. 6D where DDAs activate TRAIL/DR5 signaling through two mechanisms. First, DDAs induce ER stress that is strongly potentiated by EGFR or HER2 overexpression (Fig. 1C and (Ferreira et al., 2017)), resulting in induction of the UPR and increased DR5 expression. Previous reports have shown transcriptional upregulation of DR5 by various ER stressors (Abdelrahim et al., 2006, Tiwary et al., 2010, Xu et al., 2012, Yamaguchi & Wang, 2004, Zhang et al., 2011, Zou et al., 2008). In the case of DDAs, additional studies are required to determine which arms of the UPR are essential for activation of the DR5 promoter. TcyDTDO or RBF3 (Ferreira et al., 2015) upregulation of DR5 is not inhibited by a PERK kinase inhibitor (GSK2606414, (Axten, Medina et al., 2012)), even though upregulation of ATF4 and CHOP is blocked (Fig. S3A). PERK inhibition does not affect tcyDTDO upregulation of GRP78 or XBP1s (Fig. S3B), so XBP1s or ATF6 may participate in DR5 upregulation in response to tcyDTDO.

Second, DDAs act through mechanisms distinct from other ER stress inducers to stabilize steady-state DR5 protein levels and to induce DR5 multimerization. These mechanisms may explain the ability of tcyDTDO to induce cleavage of Caspase 8, Caspase 3, and PARP in the absence of TRAIL, and to potentiate the cytotoxicity of TRAIL. This is the first evidence that altering DR5 disulfide bonding favors multimerization and increased downstream signaling. These findings warrant additional studies to determine whether DDAs synergize with TRAIL to block tumor growth in animal models. Additional future studies should identify which cysteine residues mediate DR5 multimerization in tcyDTDO-treated cells. A recent report showed that deletion of the extracellular domain of DR5 permits oligomerization that is mediated by the transmembrane domain (Pan, Fu et al., 2019). This result indicates that the extracellular domain functions to prevent receptor oligomerization and downstream signaling in the absence of TRAIL. The extracellular domains of DR5 and DR4 contain seven disulfide bonds (see Fig. 2F) that mediate their proper folding. We speculate that DDAs alter the patterns of DR5 and DR4 disulfide bonding to allow their oligomerization and downstream signaling in the absence of TRAIL.

Although compounds that alter DR5 disulfide bonding to promote its oligomerization and downstream signaling have not previously been reported, a precedent supports such a mechanism of action for DDA-like compounds. In our previous screens of DDAs and DDA-like analogs, we observed that the compound NSC634151 was not cytotoxic to breast cancer cells (Ferreira et al., 2015). Rice and colleagues identified NSC634151 as a compound capable of blocking multiple phases of Human Immunodeficiency Virus (HIV) replication by inactivating the HIV-encoded zinc finger protein NCp7 (Rice, Baker et al., 1997). NSC634151 reacted with one or more cysteine residues in NCp7 resulting in expulsion of bound zinc. This was associated with the formation of high molecular weight, disulfide-bonded NCp7 oligomers that were only apparent on non-reducing gels. Thus, although the proteins involved are quite different, our results and those of Rice and colleagues demonstrate how thiol-reactive compounds can liberate Cys residues (that normally participate in zinc chelation in the case of NCp7, or disulfide bonds in the case of DR5) to generate inter-molecular disulfide bonds. Additional work is needed to identify the precise changes in DR5 disulfide bonding induced by DDAs.

An unexpected and striking finding is the apparent selectivity of DDAs against cancer cells over normal cells *in vitro* and *in vivo* (herein (Fig. 6C) and elsewhere (Ferreira et al., 2015, Ferreira et al., 2017)). Multiple mechanisms explain the oncotoxicity of DDAs. First, DDAs selectively induce ER stress, with associated DR5 upregulation, in the context of EGFR or HER2 overexpression (Fig. 1C). Second, breast cancer cells frequently overexpress MYC, which strongly enhances induction of apoptosis through the TRAIL/DR5 pathway (Nieminen, Partanen et al., 2007, Ricci, Jin et al., 2004, Rottmann, Wang et al., 2005, Wang, Engels et al., 2004). Third, TRAIL selectively kills cancer cells without affecting nontransformed cells (Ashkenazi et al., 1999, Griffith et al., 1999, Rieger et al., 1998, Wiley, Schooley et al., 1995). Interestingly, HCI-012 sublines selected for Lapatinib resistance exhibited higher basal EGFR and HER2 expression, and Lapatinib treatment of these lines further elevated EGFR and HER2 levels. In addition, the resistant lines showed higher MYC levels. These observations may explain why cellular responses that confer resistance to Lapatinib do not cause DDA resistance.

In summary, these findings show that DDAs exhibit unique mechanisms of anticancer action, and that they may be particularly useful against drug-resistant EGFR+ or HER2+ tumors. Studies are ongoing to determine whether DDAs potentiate the anticancer efficacy of TRAIL *in vivo* through upregulation of DR5. Efforts are also directed toward pinpointing the cysteine residues that mediate DR5 oligomerization in response to DDAs. TRAIL plays an important role in cancer cell killing by subsets of T cells and Natural Killer (NK) cells (Chen, Dieli et al., 2013, Dokouhaki, Schuh et al., 2013, Dorr, Waiczies et al., 2002, Zamai, Ahmad et al., 1998). TRAIL also can induce disruption of tumor vasculature via DR5 that is selectively expressed in tumor-associated vascular endothelial cells (Wilson, Yang et al., 2012). Therefore, it will be important in future studies to determine if DDA anticancer actions are mediated through potentiation of the actions of TRAIL that is produced by immune cells or via the induction of tumor vascular disruption.

## Materials and Methods

### Cell Culture, Preparation of Cell Extracts, and Immunoblot Analysis

The cell lines MCF10A, MDA-MB-468, BT474, T47D, SW480, and DU145 were purchased from American Type Culture Collection (ATCC) (Manassas, VA). The HCI-012 cell line was derived from a HER2+ Patient-Derived Xenograft that was originally isolated from a patient as detailed previously (DeRose et al., 2011, Ferreira et al., 2017). MCF10A cells were cultured as described previously (Jahn, Corsino et al., 2013). Unless otherwise indicated, cancer cell lines were grown in Dulbecco’s modified Eagle’s medium (GE Healthcare Life Sciences Logan, UT) supplemented with 10% fetal bovine serum (10% FBS-DMEM) in a humidified 37 °C incubator with 5% CO_2_. Cell lysates were prepared as described previously (Law, Chytil et al., 2002). Immunoblot analysis was performed employing the following antibodies purchased from Cell Signaling Technology (Beverly, MA) [Akt, #4691; P-Akt[T308], #13038; P-Akt[S473], #9271; ATF4, #11815; EGFR, #4267; HER2, #2165; HER3, #4754; IRE1, #3294; XBP1s, #12782; PARP, #9532; PERK, #5683; GRP78, #3177; CHOP, #2895; DR5, #8074; DR4, #42533; PDK1, #5662; Cleaved Caspase 8, #9496; Cleaved Caspase 3, #9664; MET, #3127; P-ERK, #9101, Rictor, #2140; MLKL, #14993; P-MLKL, #91689; PDI, #3501] and Santa Cruz Biotechnology (Santa Cruz, CA) [IGF1R, sc-713; MYC, sc-764; ERK, sc-93; Actin, sc-1616-R]. P-IRE1[Ser724] (nb100-2323ss) antibody was from Novus Biologicals.

The following reagents were purchased from the indicated sources: Tunicamycin, 2-Deoxyglucose: Sigma-Aldrich (St. Louis, MO); 2-Aminoethoxydiphenyl borate (2-APB): StressMarq Biosciences (Cadboro Bay, Victoria, Canada); Thapsigargin: AdipoGen (San Diego, CA); Cycloheximide, Rapamycin: EMD Biosciences (Darmstadt, Germany); Lapatinib: Selleck Chemicals (Houston, TX); Doxycycline, Enzo Life Science (Farmingdale, NY); CCF642, TORIN1, dithiothreitol (DTT): TOCRIS Bioscience (Minneapolis, MN); Cyclosporin A (CsA): Bioryt (Atlanta, GA); LOC14, PERK Inhibitor I (GSK2606414): Calbiochem (Burlington, Massachusetts); Q-VD-OPH, Kifunensine: Cayman Chemical (Ann Arbor, MI); *N*-ethylmaleimide (NEM): Thermo Fisher Scientific (Grand Island, NY); MG132: InvivoGen (San Diego, CA); b-AP15: MedKoo Biosciences (Chapel Hill, NC); TRAIL: PEPROTECH (Rocky Hill, NJ).

### Construction of Stable Cell Lines Using Recombinant Retroviruses and Lentiviral shRNAs

Stable cell lines were constructed as follows: Retroviral vectors encoding EGFR (Plasmid 11011 (Greulich, Chen et al., 2005)) and HER2 (Plasmid 40978 (Greulich, Kaplan et al., 2012)) were purchased from Addgene (Cambridge, MA). Stable MCF10A cell lines ectopically expressing EGFR, MYC or HER2 were prepared according to the methods used in a previous report (Law et al., 2002). MYC Plasmid 17758 was obtained from Addgene (Dai, Whitesell et al., 2007) to allow coexpression of EGFR and MYC using Puromycin and Zeocin selection, respectively.

T47D/Vector, T47D/EGFR, T47D/HER2, and T47D/EGFR/HER2 cell line construction was described in our previous report (Law, Corsino et al., 2013, Law et al., 2016). The HCI-012/LVM cells were isolated from mouse liver metastases using the conditional cell reprogramming approach of Schlegel and colleagues (Liu, Ory et al., 2012) as detailed in a previous report (Ferreira et al., 2017). Lapatinib resistant sublines 012/LVM/LR5 and 012/LVM/LR10 were developed by growing HCI-012/LVM cells in the continuous presence of either 5 µM or 10 µM Lapatinib. Lentiviral DR5 shRNA constructs were from the TRC Lentiviral shRNA Libraries from Thermo Scientific. Labeling cells with Firefly Luciferase (fLuc) was performed using Addgene Plasmid 47553 (Kang, Jensen et al., 2013). The protocol employed for developing stable cell lines with Lentiviral shRNAs was from the Thermo Scientific website: https://www.thermofisher.com/us/en/home/references/gibco-cell-culture-basics/transfection-basics.

### CRISPR/Cas9-Mediated Genome Editing

Four guide RNA sequences were designed to target the human DR5 gene (GenBank accession number: AF012628) using the online CRISPR Design tool at http://crispr.mit.edu. Sequences of the four DR5 oligonucleotide pairs are as follows: 5’-CACCGAGAACGCCCCGGCCGCTTCG-3’ and 5’-AAACCGAAGCGGCCGGGGCGTTCT-3’; 5’-CACCGCCTTGTGCTCGTTGTCGCCG-3’ and 5’-AAACCGGCGACAACGAGCACAAGGC-3’; 5’-CACCGCGCGGCGACAACGAGCACAA-3’ and 5’-AAACTTGTGCTCGTTGTCGCCGCGC-3’; and 5’-CACCGTTCCGGGCCCCCGAAGCGGC-3’ and 5’-AAACGCCGCTTCGGGGGCCCGGAAC-3’. Oligonucleotide pairs were annealed, phosphorylated with polynucleotide kinase, and cloned into the BsmBI site of LentiCRISPRv2 (Addgene #52961). Cloning of gRNA sequences into LentiCRISPRv2 was verified by sequence analysis. Viruses expressing the four different DR5 directed gRNAs were packaged using HEK 293T cells. To produce stable cell lines, target cell lines were subsequently infected with lentivirus and selected with 5 µg/ml Puromycin, as described previously (Law et al., 2002, Law et al., 2013). Clonal knockout cell lines were isolated by limiting dilution and characterized by immunoblot analysis.

### RT-PCR and RT-qPCR

Total RNA was extracted with Trizol Reagent (Invitrogen #15596-018) according to the manufacturer’s protocol. Total cellular RNA was used to synthesize first-strand cDNA by using the PCR conditions listed: 25 C for 10 min, 42 C for 30 min, and 95 C for 5 min. PCR was subsequently performed using either DR5 or β-Actin primers. The primer sequences for DR5 are 5’-TCCACCTGGACACCATATCTCAGAA-3’ and 5’-TCCACTTCACCTGAATCACACCTG-3’ and the primer sequences for β-Actin are 5’-GGATGCAGAAGGAGATCAC-3’ and 5’-AAGGTGGACAGCGAGGCCAG-3’. The PCR products were visualized on 3% agarose gels with ethidium bromide staining under UV transillumination with a digital camera system, and quantified using NIH ImageJ. Real time-qPCR was performed with the QuantStudio 6 Flex system (ThermoFisher #4485691) using PowerUp from Invitrogen (https://www.thermofisher.com/order/catalog/product/A25742). The expression level of DR5 was normalized to β-Actin, and the cycle threshold value of the sample was used to calculated the relative gene expression level = 2^-(Ct target-Ct actin)^. The relative change in gene expression compared with the control group was expressed as fold change, and calculated by the 2^-ΔΔCt^ method (Schmittgen & Livak, 2008).

### Protein Synthesis Assays

Protein synthesis assays were carried out as detailed previously (Law, Waltner-Law et al., 2000) using ^3^H-Leucine (cat. # NET460001MC) obtained from Perkin Elmer (Waltham, MA).

### Luciferase Transcriptional Reporter Assays

Reporter assays were performed as described previously (Ferreira et al., 2017) using the DR5 reporter construct obtained from Addgene (Plasmid 16012 (Takimoto & El-Deiry, 2000)).

### Cell Viability Assays

Cell viability was evaluated using the MTT (3-(4,5-dimethylthiazol-2-yl)-2,5-diphenyltetrazolium bromide) assay. MTT assays were carried out according to the Manufacturer’s instructions (kit CGD1, Sigma-Aldrich, St. Louis, MO).

### In Vivo Tumor Studies, Histochemical, and Immunohistochemical Analysis

Mice were housed, maintained, and treated in the Animal Care Service Center at the University of Florida in accordance with Institutional Animal Care and Use Committee (protocol number: 201608029). NOD-SCID-G) mice were obtained from Jackson Laboratories (Bar Harbor, ME). Breast cancer liver metastasis was initiated by injecting 1 × 10^6^ cancer cells into the #4 mammary fat pads, or 200,000 cells into the lateral tail veins of adult female NSG mice.

Tissue samples were fixed in 4% paraformaldehyde/PBS and paraffin embedded. Tissue sectioning, hematoxylin and eosin (H&E) staining, and immunohistochemical staining for cleaved Caspase 3 (Cat. #9664. Cell Signaling Technology) were performed by the University of Florida Molecular Pathology Core (https://molecular.pathology.ufl.edu/).

### Statistics

In Fig. 6A data obtained from The Cancer Genome Atlas (TCGA; https://tcga.xenahubs.net/download/TCGA.BRCA.sampleMap/HiSeqV2_PANCAN.gz) [dataset ID: TCGA_BRCA_exp_HiSeqV2_PANCAN] were used to examine the relationships between tumor expression of EGFR and MYC at the mRNA level and patient survival. Gene expression data for 1176 patients with invasive breast carcinoma was measured by RNAseq and mean-normalized across all TCGA cohorts. EGFR and MYC were the two genes of interest in our study. Patients ranked based on the expression of EGFR and MYC were classified into four groups, named “low EGFR low MYC (N=359)”, “low EGFR high MYC (N=220)”, “high EGFR low MYC (N=225)”, and “high EGFR high MYC (N=372)”. Overall Survival (OS) was compared among these groups. The F test was used to compare the variance between groups (*P* > 0.05, ns; *P* ≤ 0.05, *; *P* ≤ 0.01, **; *P* ≤ 0.001, ***).

Statistical analyses for synergies between drugs were performed using CalcuSyn software (http://www.biosoft.com/w/calcusyn.htm). The combination index (CI) was calculated by applying the Chou-Talalay method and was used for synergy quantification (Chou, 2010). Student’s *t*-test was used for comparisons in both *in vitro* and *in vivo* experiments. All *P* values are two-tailed, and both *P* values and statistical tests are mentioned in either figures or legends.

### Chemical syntheses of DDAs

RBF3 and tcyDTDO were prepared as described previously (Ferreira et al., 2015, Wang et al., 2019).

## Acknowledgements

These studies were supported in part by grants from the Florida Breast Cancer Foundation (BL and RC), the Ocala Royal Dames for Cancer Research (BL), the Florida Department of Health Bankhead-Coley Program (4BF03, BL), the Collaboration of Scientists for Critical Research in Biomedicine (CSCRB, Inc., BL), and a UF-Health Cancer Center pilot project award (BL, RC, and CH). This work was supported in part by the Office of the Assistant Secretary of Defense for Health Affairs through the Breast Cancer Research Program under Award Nos. W81XWH-15-1-0199 (BL) and W81XWH-15-1-0200 (RC). RBF is grateful to the University of Florida for a Graduate School Fellowship. Finally, we thank breast cancer research advocate Mrs. Jeri Francoeur for her advice and support.

## Author contributions

Mengxiong Wang performed most of the experiments, including the cell culture, immunoblot analyses, CRISPR/Cas9-mediated genome editing, cell viability assays, animal studies, and data analyses. Mary E. Law contributed to the construction of stable cell lines using recombinant retroviruses and lentiviral shRNAs, and isolated total RNA from cells. Bradley J. Davis performed luciferase transcriptional reporter assays and contributed to the animal studies. Elham Yaaghubi, Amanda F. Ghilardi, Renan B. Ferreira, and Ronald K. Castellano designed and synthesized DDAs. Chi-Wu Chiang, Olga A. Guryanova, and Daniel Kopinke provided reagents and reviewed the manuscript. Coy D. Heldermon provided breast cancer patient samples. Brian K. Law designed the studies and wrote the manuscript with Mengxiong Wang.

## The paper explained

### Problem

Treatment options for breast tumors that have disseminated and acquired resistance to current therapeutics are limited. Novel treatment strategies that are effective against drug-resistant, metastatic cancer are critically needed.

## Results

Herein, we have demonstrated that a novel chemical class of anti-cancer agents, termed DDAs, exhibit strong anticancer activity by stimulating Death Receptor 5 (DR5)-mediated breast cancer cell death. When activated by its ligand, TRAIL, DR5 induces cancer cell death. Our results indicate that DDAs are unique in causing DR5 accumulation and oligomerization through mechanisms involving altered DR5 disulfide bonding. DDAs prominently kill both the primary tumor and tumor metastasis in our mouse model without affecting surrounding normal tissues, and synergize with TRAIL to kill drug-resistant breast cancer cells.

### Impact

This study demonstrates that DDAs may be a promising therapeutic strategy for patients with highly resistant and metastatic breast cancer.

