## Supplemental Material for "Mechanistic Elucidation of the Antitumor Properties of a Novel Death Receptor 5 Activator"

### Wang et al. - Appendix

**Appendix Figure Legends**

**Appendix Figure S1: DDAs increase DR5 mRNA.** A. Luciferase transcriptional reporter assays were performed in HEK 293 cells using a DR5 transcriptional reporter construct. Cells were treated with DMSO vehicle or 5 μM tcyDTDO for 24 h prior to luciferase assays. B. RT-PCR of DR5 was performed on the MDA-MB-468 cell line with indicated treatments. DR5 mRNA levels were measured by NIH ImageJ analysis and were normalized to corresponding β-Actin mRNA levels. C., D. Immunoblot analysis of MDA-MB-468 and SKBR3 cells engineered to express CHOP in a tetracycline-inducible manner (468/tet-CHOP) and the corresponding control cell line (468/tet-Puro) after the indicated 24 h treatments. E. Immunoblot analysis of the indicated stable BT474 shRNA-mediated knockdown cell lines after 24 h treatment with vehicle or 20 □M RBF3. Error bars in S1A and S1B represent standard deviation.

**Appendix Figure S2: TcyDTDO-induced DR5 electrophoretic mobility shifts.** A. MDA-MB-468 cells were treated for 24 h with 12.5 ng/ml TRAIL and the indicated concentrations of the other agents, and extracted with boiling 2X SDS-PAGE sample buffer containing 100 mM NEM. Samples were then sonicated, clarified by centrifugation, and analyzed by immunoblot. B. BT474 cells engineered to express DR5 in a doxycycline-inducible manner (BT474-tet-DR5) were treated for 24 h with or without 1 μg/ml doxycycline in the presence or absence of 5 μM tcyDTDO, 100 nM Thapsigargin, 250 ng/ml Tunicamycin, 3 mM DTT, 15 μM CsA, or 12.5 ng/ml TRAIL as indicated. Cells were extracted and analyzed as described in Fig. S2A. C. MDA-MB-468 cells were treated for 24 h as indicated and analyzed by immunoblot as described in Fig. S2A. In A-C, blue arrows indicate bands corresponding to the long and short forms of DR5 under control conditions and red arrowheads show bands corresponding to DR5 migration after treatment with tcyDTDO. Note that tcyDTDO reduces the migration of both DR5 isoforms. D. Micrographs of 012/LVM/fLuc cells treated for 24 h with vehicle (DMSO), 10 μM tcyDTDO, 25 ng/ml TRAIL, or tcyDTDO + TRAIL. E. Micrographs showing the morphology of BT474 cells after 24 h of treatment as indicated.

**Appendix Figure S3: TcyDTDO upregulation of DR5 is not blocked by PERK inhibition.** A. MDA-MB-468 (left panel) or BT474 cells (right panel) were treated for 24 h as indicated and subjected to immunoblot analysis. B. 012/LVM/LR10 cells were treated for 24 h as indicated and subjected to immunoblot analysis.

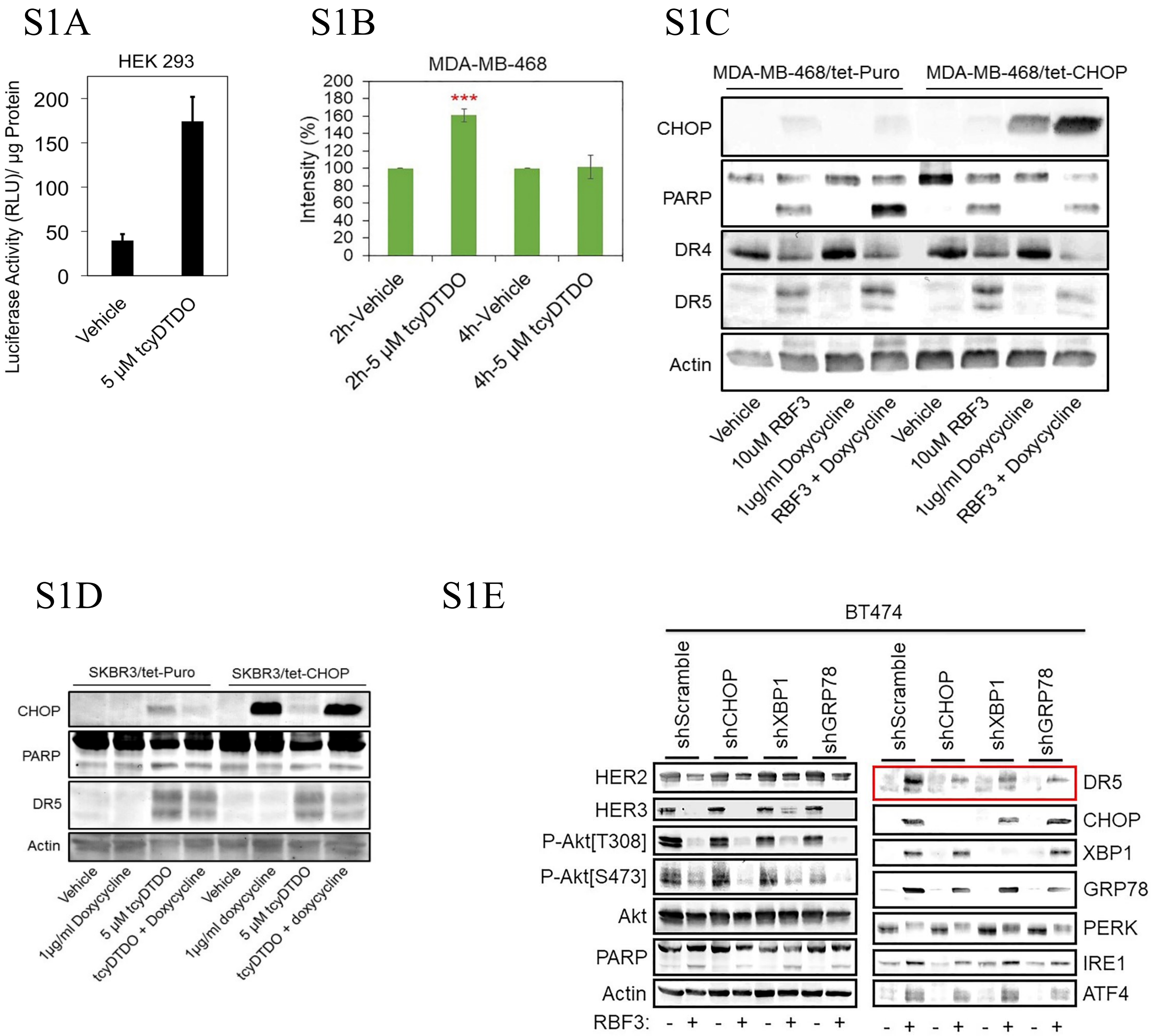

*Appendix Figure S1*

S2A

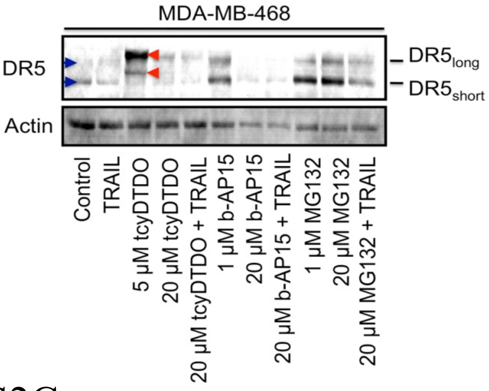

S2B

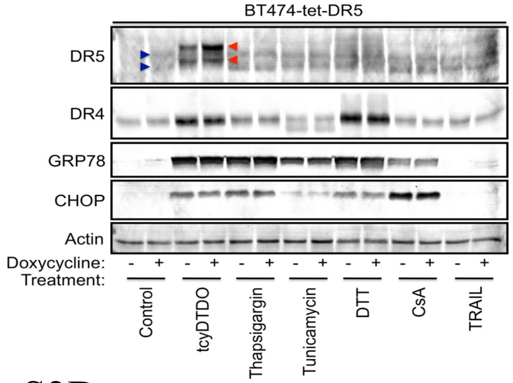

S2C

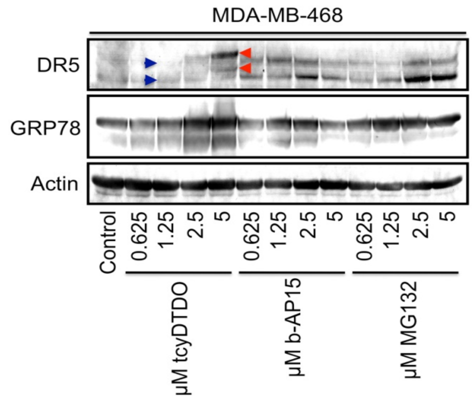

S2D

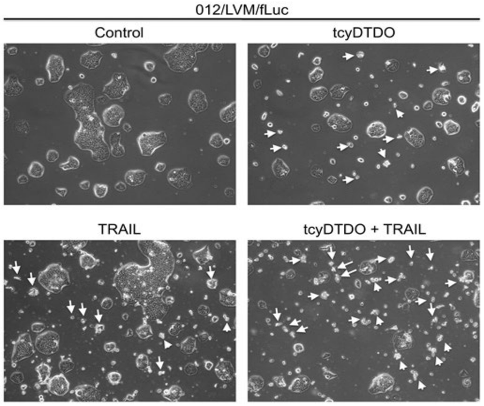

S2E

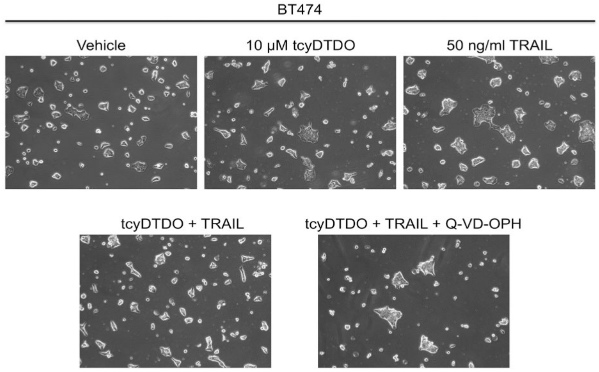

*Appendix Figure S2*

S3A

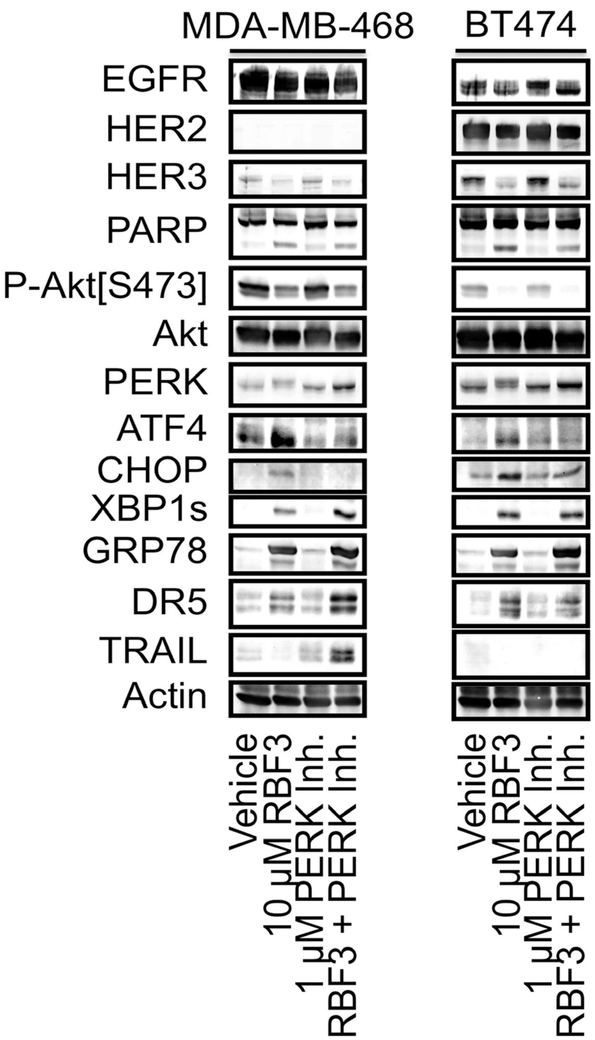

S3B

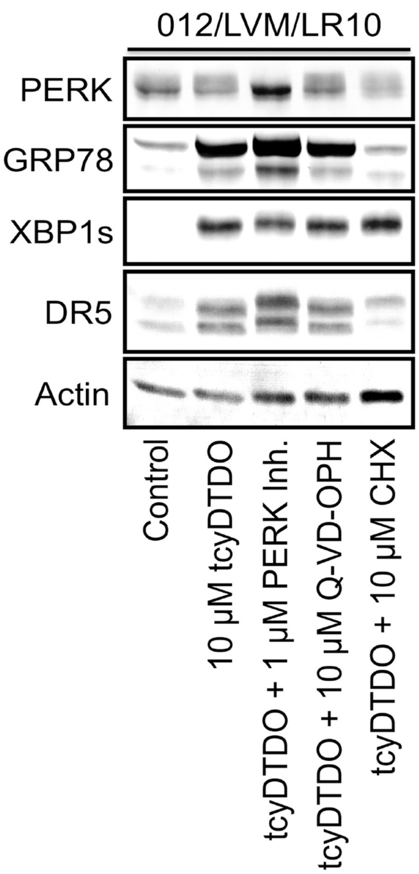

*Appendix Figure S3*
